# An *OpIE2*-DsRed marker disrupts female blood-feeding and shortens lifespan in the malaria vector *Anopheles gambiae*

**DOI:** 10.1101/2025.10.14.682374

**Authors:** Arad Sarig, Evyatar Sar-Shalom, Ebrima SM Kolley, Elad Shmuel Yonah, Lee Benjamin Lamdan, Ariel Roselin Lewin, Tamir Partosh, Gur Pines, Jonathan D. Bohbot, Philippos Aris Papathanos

**Affiliations:** Department of Entomology, Institute of Environment Sciences, Robert H. Smith Faculty of Agriculture, Food and Environment, Hebrew University of Jerusalem, Rehovot, Israel; Institute of Plant Protection, Agricultural Research Organization - Volcani Institute, Rishon LeZion, Israel

## Abstract

*Anopheles gambiae* is one of the principal vectors of human malaria. Over the past two decades, transgenic mosquito strains have been essential tools for studying mosquito biology and developing genetic control strategies such as gene drives. Mosquito transformants are typically identified using fluorescent markers, which are assumed to be phenotypically neutral. While generating CRISPR-based gene drive strains carrying an *OpIE2*-DsRed marker we unexpectedly found that transgenic females were unable to blood-feed and were consequently sterile, whereas males initially appeared normal and fertile. Given the potential utility of dominant, female-specific sterility for mosquito control, we established additional strains controlling for transgene content and integration site, confirming that the *OpIE2*-DsRed cassette caused the defect. Behavioral assays showed that females exhibited normal attraction to a membrane feeder but failed to initiate blood-feeding, performing repeated cycles of probing and proboscis grooming in rapid succession before ultimately leaving the feeder unfed. Microscopy showed that both sexes possessed a distally curved proboscis, providing a morphological explanation for the blood-feeding defect of females and the reduced male lifespan. A second promoter variant (*OpIE2b*), differing in flanking sequences at the IE-2 junction, drove strong marker expression without impairing blood-feeding or longevity. These findings demonstrate that minor differences in promoter architecture can produce major, unexpected phenotypic effects. *OpIE2b* provides a robust, phenotypically neutral marker for *An. gambiae* research, while *OpIE2a* highlights the need for rigorous validation of transgenic components intended for research and applied releases.

## Introduction

The African malaria mosquito *Anopheles gambiae* is among the most efficient vectors of *Plasmodium* parasites and remains a major public-health challenge in sub-Saharan Africa. Transmission occurs exclusively through the blood-feeding of adult females, which digest blood to obtain nutrients required for oocyte development and egg production. This link between blood-feeding and reproduction makes female feeding behavior a critical determinant of both the vectorial capacity and mosquito population dynamics.

Since stable germline transformation of *Anopheles* mosquitoes was first demonstrated in the early 2000s (Catteruccia et al. 2000; Grossman et al. 2001), transgenic strains have become important tools for mosquito research. They now underpin a wide range of studies in mosquito biology, from malaria transmission biology (Pascini et al. 2022) to olfaction and sensory behavior (Georgiades et al. 2023; Giraldo et al. 2024; Maguire et al. 2022; Riabinina et al. 2016) and insecticide resistance (Adolfi et al. 2019). Transgenic approaches are also integral to the development of genetic control strategies, including gene drives designed for population suppression and replacement (Carballar-Lejarazú et al. 2020; Haber et al. 2024; Kyrou et al. 2018; Li et al. 2025; Strampelli et al. 2025; Tolosana et al. 2025; Verkuijl et al. 2025).

These advances depend not only on the ability to introduce and stably integrate transgenes into the mosquito genome, but also on reliably identifying transformed individuals. In practice, this is achieved through fluorescent reporters driven by 5’ and 3’ regulatory sequences (hereafter 5’ regulatory sequences are referred to as promoters for simplicity) that ensure strong and easily detectable expression, most often during larval stages when screening is performed under a fluorescent microscope. An ideal reporter is bright throughout development, spectrally distinct from other markers (e.g., eGFP vs DsRed), and compatible with multiple constructs, for instance through tissue-specific expression patterns (e.g., eye-versus muscle-specific). Differences in brightness between homozygotes and heterozygotes can sometimes aid husbandry and genetics, though these are highly dependent on the genome integration site and are more reliably distinguished using automated optical sorters (e.g., COPAS) than by eye (Marois 2024). Above all, reporters should be phenotypically neutral, imposing no measurable fitness costs that might confound experiments or limit their application.

Several promoters have been used to drive fluorescent marker expression in *Anopheles gambiae*, including the synthetic *3xP3* promoter, which drives strong eye- and neuron-specific fluorescence (Berghammer, Klingler, and Wimmer 1999) and the heterologous *Actin5C* promoter from *Drosophila melanogaster*, which provides broad somatic expression (Volohonsky et al. 2015). Endogenous promoters have also been developed, including the *polyubiquitin* promoter (Adolfi et al. 2018) and the α*-tubulin-1b* promoter (Lycett, Amenya, and Lynd 2012) which drive multi-tissue expression. Viral regulatory elements like *hr5-IE1* have also been used to drive ubiquitous expression (Grossman et al. 2001; Meredith et al. 2013). The *OpIE2* promoter, derived from the immediate early gene 2 (*IE-2*) of the *Orgyia pseudotsugata* multicapsid nucleopolyhedrovirus (OpMNPV) (Theilmann and Stewart 1992) has proven highly effective in diverse insect systems. The *OpIE2* promoter works well in lepidopteran and dipteran cell line transfections (Pfeifer et al. 1997; Xu et al. 2013) and in transgenic strains of several insect species, including in *Drosophila melanogaster* (Buchman et al. 2018; Kandul et al. 2019), *Drosophila suzuki* (Liu et al. 2025; Witherbee and Gamez 2025), *Bombyx mori* (Li et al. 2015; Tang et al. 2018), *Pectinophora gossypiella* (Simmons et al. 2011)*, Plutella xylostella* (Martins et al. 2012)*, Ceratitis Capitata* (Davydova et al. 2023) and *Aedes aegypti* (Estevez-Castro et al. 2024; Li et al. 2017; Rouyar et al. 2024; Vainer et al. 2024) *Aedes albopictus* (Lutrat et al. 2022; Zaada et al. 2025) and *Culex quinquefasciatus* (Feng et al. 2021). However, its performance in *An. gambiae* has not been systematically examined.

Here, we report that expression of DsRed in *An. gambiae* under a specific variant of the *OpIE2* promoter severely impairs female blood-feeding and reduces male longevity. To investigate this effect, we generated a number of strains controlling for genomic integration site and construct elements, characterized female feeding behavior, and tested an alternative *OpIE2* promoter sequence. The alternative promoter maintained marker visibility while restoring normal feeding, underscoring that promoter choice can have profound effects on mosquito biology and is critical for both transgenic tool design and genetic control strategies.

## Results

### Generation of *OpIE2-*DsRed transgenic strains

In previous work, we used an *OpIE2*-DsRed fluorescent marker in *D. melanogaster* strains containing split Cas9 and gRNA sex ratio distorters (Haber et al. 2024). Seeking to develop similar capacities in *An. gambiae*, we cloned this *OpIE2*-DsRed to label sgRNA transgenic constructs, complementing available Cas9 strains marked by fluorescent proteins driven by the neuron-specific *3xP3* promoter. Four transformation constructs were initially generated, each containing *OpIE2*-DsRed and single *U6*-driven sgRNA targeting different sex chromosome sequences (**Figure 1A**). The constructs included *piggyBac* repeats for transposase-mediated integration and an *attB* site for *phiC31*-mediated site-specific recombination. Constructs were injected either into wild type eggs with a *piggyBac* transposase helper or into eggs of the *attP*-E strain, which carries a well-characterized, *3xP3*-CFP-labelled *attP* docking site on chromosome 3 (Meredith et al. 2011). In total, eight independent strains were generated, four with random *piggyBac* insertion (construct #1) and three in the *attP*-E line (constructs #2-4; **Supplementary Table 1**).

**Figure 1.**
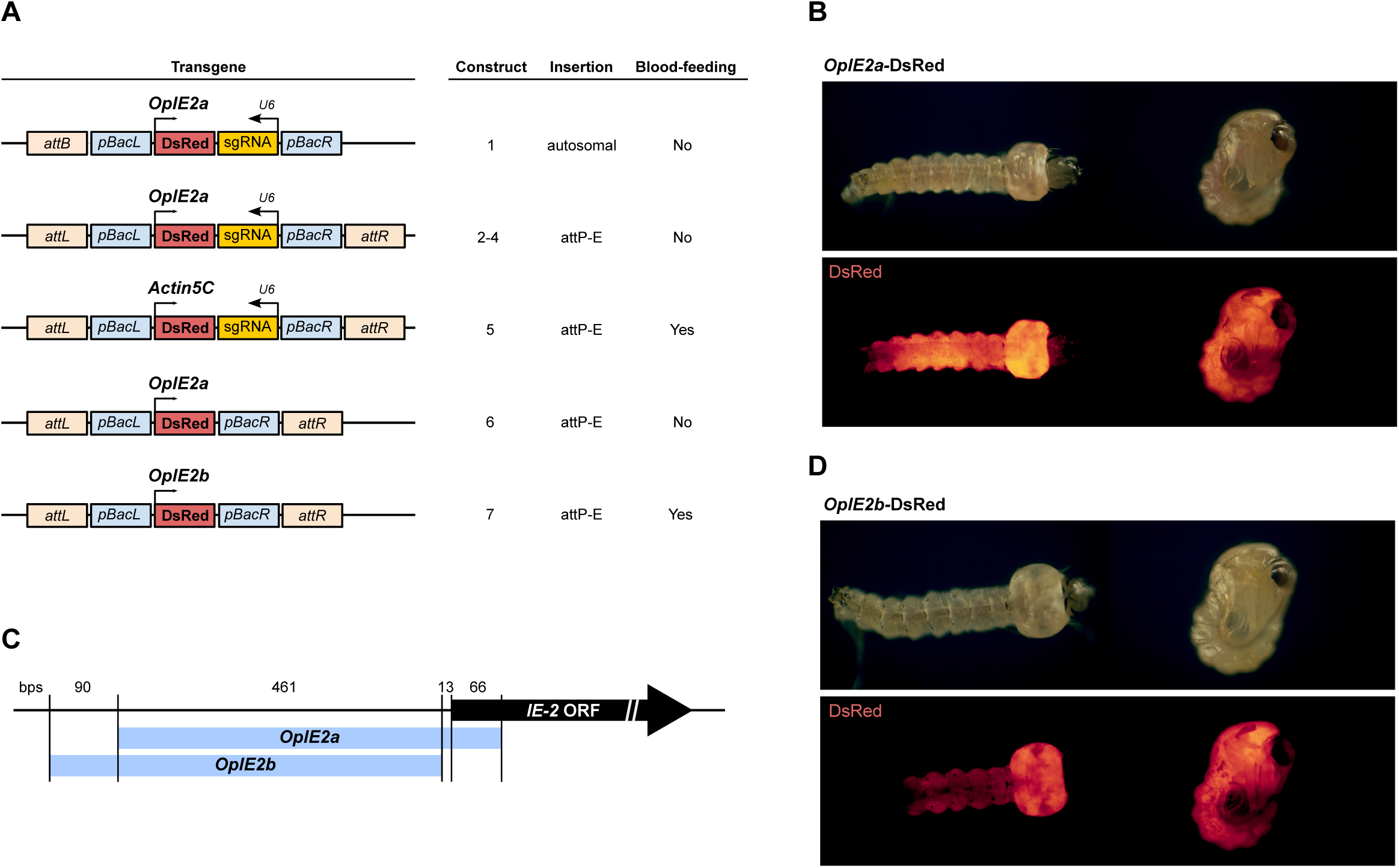
Transgenic constructs and marker expression in *OpIE2a* and *OpIE2b* promoters driving DsRed. **(A)** Transgenic construct design carrying either *OpIE2a*, *OpIE2b*, or *Actin5C* promoters driving DsRed, with or without a *U6*-sgRNA cassette. All were flanked by *piggyBac* inverted repeats and *attB* recombination sites enabling *phiC31*-mediated site-specific integration. The genomic location and female blood-feeding phenotype are summarized on the right. **(B)** Brightfield and fluorescence of fourth-instar larvae and pupae expressing DsRed under the *OpIE2a* promoter. **(C)** Alignment of the *OpIE2a* and *OpIE2b* promoter sequences relative to the *Orgyia pseudotsugata* immediate early-2 (*IE-2*) gene. Both promoters share a common 461 bp core region, but *OpIE2a* extends 66 bp into the *IE-2* coding sequence, while *OpIE2b* includes an additional 90 bp upstream region and terminates before the start codon. **(D)** Brightfield and fluorescence of fourth-instar larvae and pupae expressing DsRed under the *OpIE2b* promoter.

Larvae of all strains displayed bright DsRed fluorescence in thoracic and abdominal segments, and weaker expression in the larval head (**Figure 1B**). In pupae, DsRed fluorescence was bright throughout the entire body (**Figure 1B**). However, while establishing the strains, we observed that adult transgenic females from all eight lines failed to blood-feed (**Figure 1A**), and oviposition bowls provided 48 hours later contained no eggs - a phenotype described in detail below. In contrast, transgenic males appeared physiologically normal and fertile, producing offspring when crossed to wild-type females and enabling maintenance of the strains as heterozygotes. Because the defect was consistent across both *piggyBac* and *phiC31* integrations, we concluded that it likely resulted from the *OpIE2*-DsRed cassette itself rather than the insertion site or sgRNA sequence, which differed between constructs.

To test this, we generated a new transformation construct in which DsRed was driven by the *Actin5C* promoter instead of *OpIE2*. When integrated at the same *attP-E* site, this construct restored female blood-feeding (construct #5; **Supplementary Table 1; Figure 1A**), suggesting that the phenotype was specific to *OpIE2*-DsRed expression. We confirmed this by generating a new construct containing only the *OpIE2*-DsRed cassette and integrating it into the *attP*-E site (construct #6; **Supplementary Table 1; Figure 1A**). As this strain exhibited the same non-blood-feeding phenotype, we concluded that *OpIE2-*DsRed expression alone was sufficient to interrupt female blood-feeding.

When we analyzed the cloned *OpIE2* promoter sequence, we noticed that the promoter derived from the *Drosophila* constructs differed from another *OpIE2* promoter that we recently used in *Aedes albopictus* (Zaada *et al*., 2025). To better understand these differences we aligned both promoters, which we called *OpIE2a* (for *D. melanogaster* constructs) and *OpIE2b* (for *A. albopictus* constructs) to the *OpMNPV IE-2* gene region. Both promoters shared a 461 bp core sequence, but there were several differences in the cloned flanking sequences. (**Figure 1C**). The *OpIE2a* promoter extended 66 bp into the viral *IE-2* coding sequence, including the start codon and fusing 22 amino acids of *IE-2* to the N-terminus of DsRed. The *OpIE2b* promoter included an additional 90 bp upstream sequence, but terminated prior to the *IE-2* start codon in its 3’ end, where it connects to DsRed. The 3’ end of the *OpIE2b* promoter also contained *T7* and *EM7* expression elements and restriction sites retained from previous cloning steps.

To test whether these sequence differences accounted for the phenotype, we generated a new strain carrying the *OpIE2b*-DsRed cassette integrated into the same *attP-E* site (construct #7; **Supplementary Table 1; Figure 1A**). DsRed expression driven by *OpIE2b* was slightly weaker in larvae, pupae and adults compared to *OpIE2a* but followed a similar spatial pattern (**Figure 1C**). Importantly, *OpIE2b*-DsRed females blood-fed normally (**Figure 1A**), demonstrating that *OpIE2a*-DsRed caused the feeding defect.

### Blood-feeding capacity is impaired in *OpIE2a-DsRed* females

To characterize and quantify the blood-feeding defect, we established triplicate cages of 14-20 *OpIE2a*-DsRed or *OpIE2b*-DsRed females, previously mated to wild type males, alongside wild type control females. Females were offered a membrane feeder containing bovine blood for 25 minutes, and a camera inside each cage recorded female behavior (**Figure 2A**). At the end of the assay, all females were aspirated from their cages to record the number of blood-fed females. *OpIE2b*-DsRed females fed normally, with feeding rates indistinguishable from wild type females (*p* = 0.254, unpaired two-sided *t*-test; **Figure 2B**). In contrast, none of the *OpIE2a-* DsRed females fed during the 25-minute window (*p* = 0.0002, unpaired two-sided *t*-test; **Figure 2B**). To test whether additional stimulation could elicit feeding, *OpIE2a-*DsRed females were given a 15-minute rest and re-offered fresh blood for an additional 30 minutes, accompanied by intermittent human breath stimulation to mimic host-associated cues. Eventually a small fraction of *OpIE2a-*DsRed females managed to feed (median = 6.25%; four females across all replicates; **Figure 2B**).

**Figure 2.**
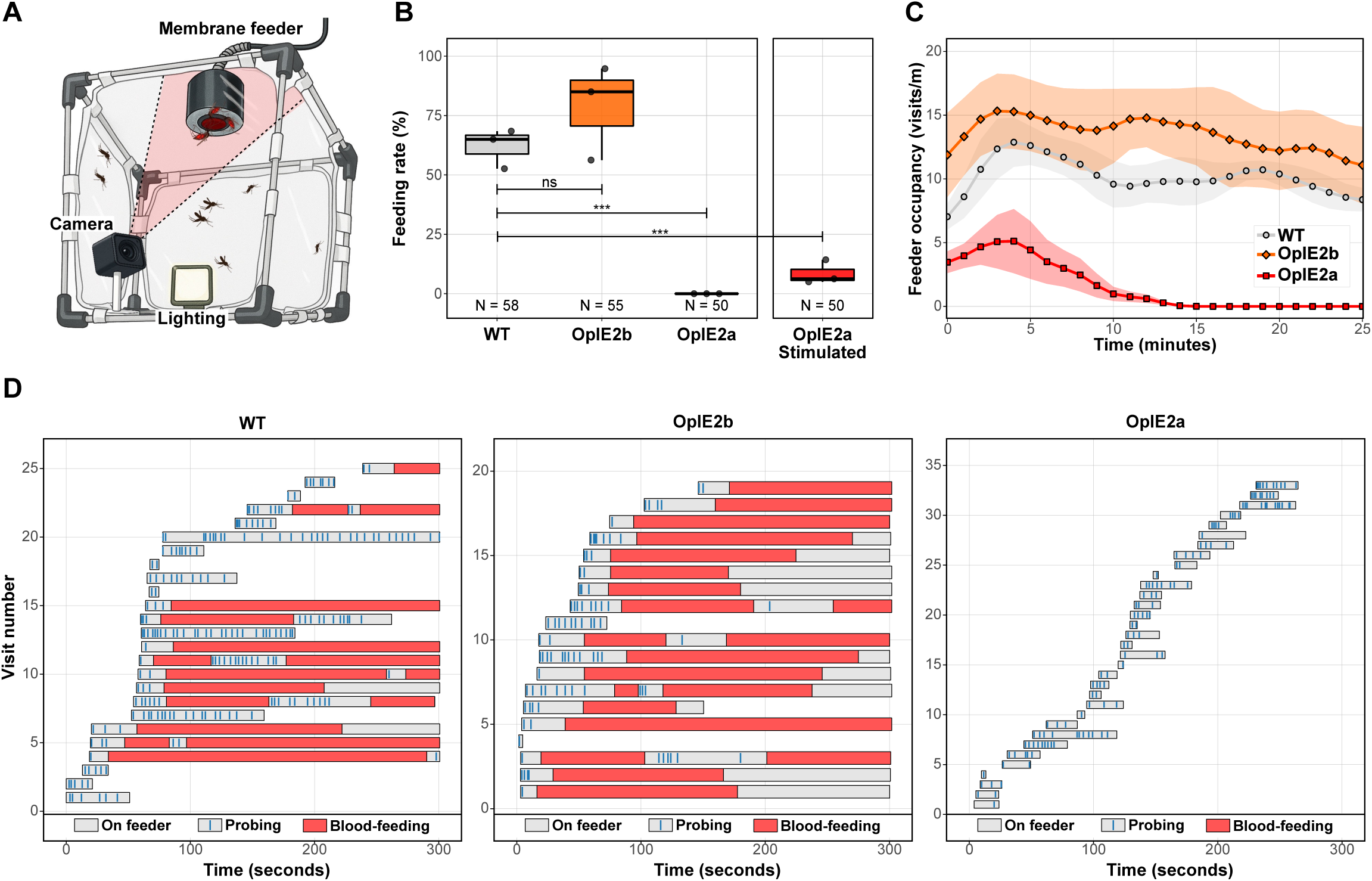
Blood-feeding behavior of *OpIE2*-DsRed females. **(A)** Illustration of the experimental setup for the blood-feeding assay. Females were placed in minicages containing a camera and light source. A membrane-feeder filled with bovine blood was placed on top of the cage at the beginning of the experiment. **(B)** Feeding success of wild type, *OpIE2b* and *OpIE2a* females. *OpIE2b*-DsRed females fed normally, whereas *OpIE2a*-DsRed females failed to feed. Re-exposure of *OpIE2a*-DsRed females to a new feeder following 15min of resting with human-breath stimulation induced only occasional feeding. Boxplots show medians ± IQR; points represent independent replicate cages. N indicates the total number of females tested per treatment. Statistical significance was derived from unpaired two-sided *t*-tests, assuming equal or unequal variances, comparing the feeding rates between the different strains to the wild-type control (αL=L0.05). **(C)** Feeder occupancy over time quantified by a YOLO-based behavioral tracking network. Solid lines show Gaussian-smoothed mean visits per minute ± SEM across (shaded ribbon) replicate cages. *OpIE2a*-DsRed females spent significantly less time on the membrane-feeder and their visits declined rapidly compared to WT and *OpIE2b*-DsRed controls. **(D)** Manually annotated behavioral ethograms during the first five minutes of the blood-feeding assay: Bars represent sequential visits by individual females, color-coded as on-feeder (gray), probing (blue), and blood-feeding (red). *OpIE2a*-DsRed females displayed repeated short probing attempts without transition to sustained feeding.

Recordings of female blood-feeding behavior were analyzed using a fine-tuned YOLOv11l-based object detection model (Khanam and Hussain, 2024) adapted for mosquito detection by Vainer et al. (2024). *OpIE2a*-DsRed females showed a significantly steeper decline in feeder visits over time compared to wild-type controls (interaction β = –0.158 ± 0.038 SE, *p* < 0.001), whereas *OpIE2b*-DsRed females did not differ significantly (β = –0.047 ± 0.038 SE, *p* = 0.21). At the beginning of the assay (time = 0), *OpIE2a*-DsRed females already exhibited fewer visits than wild type (–6.66 ± 2.23 SE, *p* = 0.0028), while *OpIE2b*-DsRed females were comparable to controls (*p* = 0.08; **Figure 2C**).

Behavioral ethograms for the first five minutes of each assay were also generated, annotating arrival, probing, blood-feeding and departure events for each female. Wild-type and *OpIE2b*-DsRed females began probing shortly after arrival and remained stationary once feeding commenced, spending significantly more time on the feeder overall (**Figure 2C; Supplementary Figure 1**). In contrast, *OpIE2a*-DsRed females displayed normal attraction to the feeder, probing the membrane at a similar frequency to wild-type females, but all failed to initiate blood-feeding. Instead, they performed repeated cycles of unsuccessful probing followed by proboscis grooming, ultimately leaving the feeder unfed after brief visits (**Figure 2C; Supplementary Figure 1**). These observations indicate that the *OpIE2a*-DsRed defect impairs feeding initiation rather than attraction.

### *OpIE2a*-DsRed adults exhibit a curved proboscis

To investigate whether the behavioral defect corresponded to morphological abnormalities, we examined the anterior feeding and sensory appendages of adult mosquitoes by stereofluorescence microscopy. All *OpIE2a*-DsRed females displayed a conspicuous distal curvature of the proboscis and maxillary palps, in contrast to the straight, parallel alignment characteristic of wild-type and *OpIE2b*-DsRed females (**Figure 3A-C; Supplementary Figure 2)**. The curvature averaged 21.3**°** along the distal end of the proboscis forming an abrupt ventral deflection. The paired maxillary palps also appeared bent at the same position as the proboscis.

**Figure 3.**
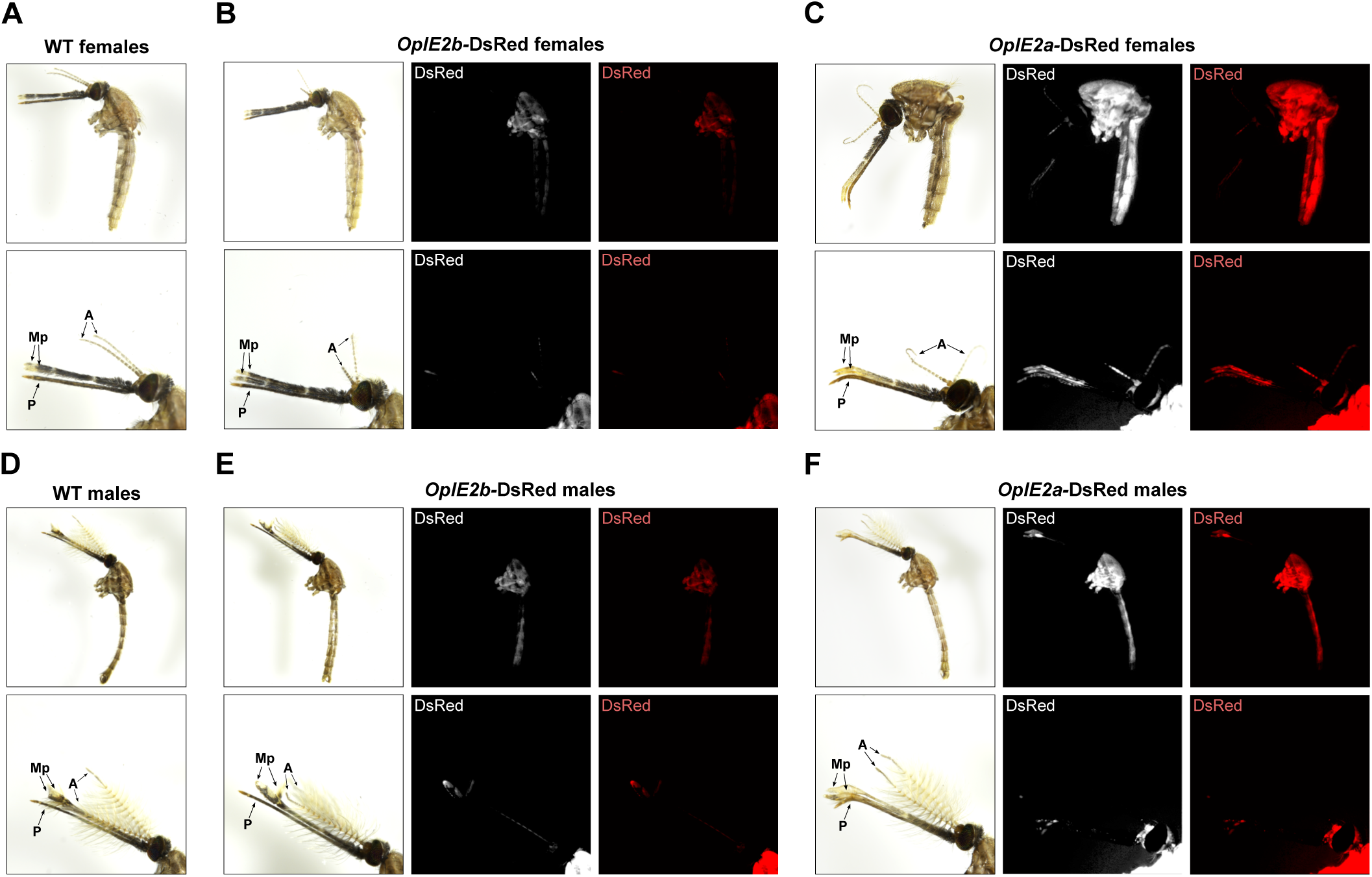
Proboscis morphology and DsRed fluorescence in *Anopheles gambiae* adults carrying *OpIE2*-driven markers. **(A–C)** Females and (**D–F)** males from wild-type (WT), *OpIE2b*-DsRed, and *OpIE2a*-DsRed strains imaged under bright-field (left) and fluorescence (middle, grayscale; right, red channel) illumination. **(A,D)** WT adults display a straight proboscis with no fluorescence. **(B,E)** *OpIE2b*-DsRed adults show strong, widespread DsRed expression but retain normal proboscis morphology. **(C,F)** *OpIE2a*-DsRed adults exhibit a pronounced distal curvature of the proboscis and strong DsRed signal localized on the proboscis and maxillary palps. Images were acquired under identical exposure settings to permit direct comparison of fluorescence intensity and anatomical deformation across strains and sexes.

Because the feeding defect initially appeared female-specific, we also examined males as controls (**Figure 3D-F)**. Unexpectedly, *OpIE2a*-DsRed males exhibited the same deformity, with curvature of both the proboscis and palps even more pronounced than in females (mean of 39.2**°**). The male proboscis often appeared twisted or bent laterally at the tip, and in severe cases the palps crossed over one another, partially covering the proboscis. In contrast, *OpIE2b*-DsRed males retained normal morphology, with straight, evenly aligned proboscis and palps indistinguishable from wild type.

Fluorescence imaging revealed notable DsRed signals within the proboscis and maxillary palps of *OpIE2a*-DsRed adults, but not in *OpIE2b*-DsRed mosquitoes imaged under identical exposure settings. The co-occurrence of proboscis-localized DsRed expression and curvature in *OpIE2a*-DsRed individuals suggests that promoter sequences influence expression in anterior appendages, coinciding with the observed morphological abnormalities.

### *OpIE2a-*DsRed males exhibit reduced adult longevity

To determine whether the *OpIE2a*-DsRed construct was associated with broader fitness costs, we assessed reproductive output and survival in both sexes. Among the few *OpIE2a*-DsRed females that successfully blood-fed after stimulation (n = 4), only two laid eggs. The number of eggs produced by these females was markedly lower than in controls, with a median of 28 eggs compared to 97.5 in wild-type and 74.5 in *OpIE2b*-DsRed females (unpaired two-sided *t*-test, *p* = 0.042; **Figure 4A**). In contrast, egg hatching rates did not differ significantly among strains (**Figure 4B**), indicating that fertilization and embryonic development were unaffected. When the blood-feeding assay was repeated with daughters of these females, none successfully blood-fed, confirming that the phenotype was heritable and fully penetrant (**Supplementary Table 2**).

**Figure 4.**
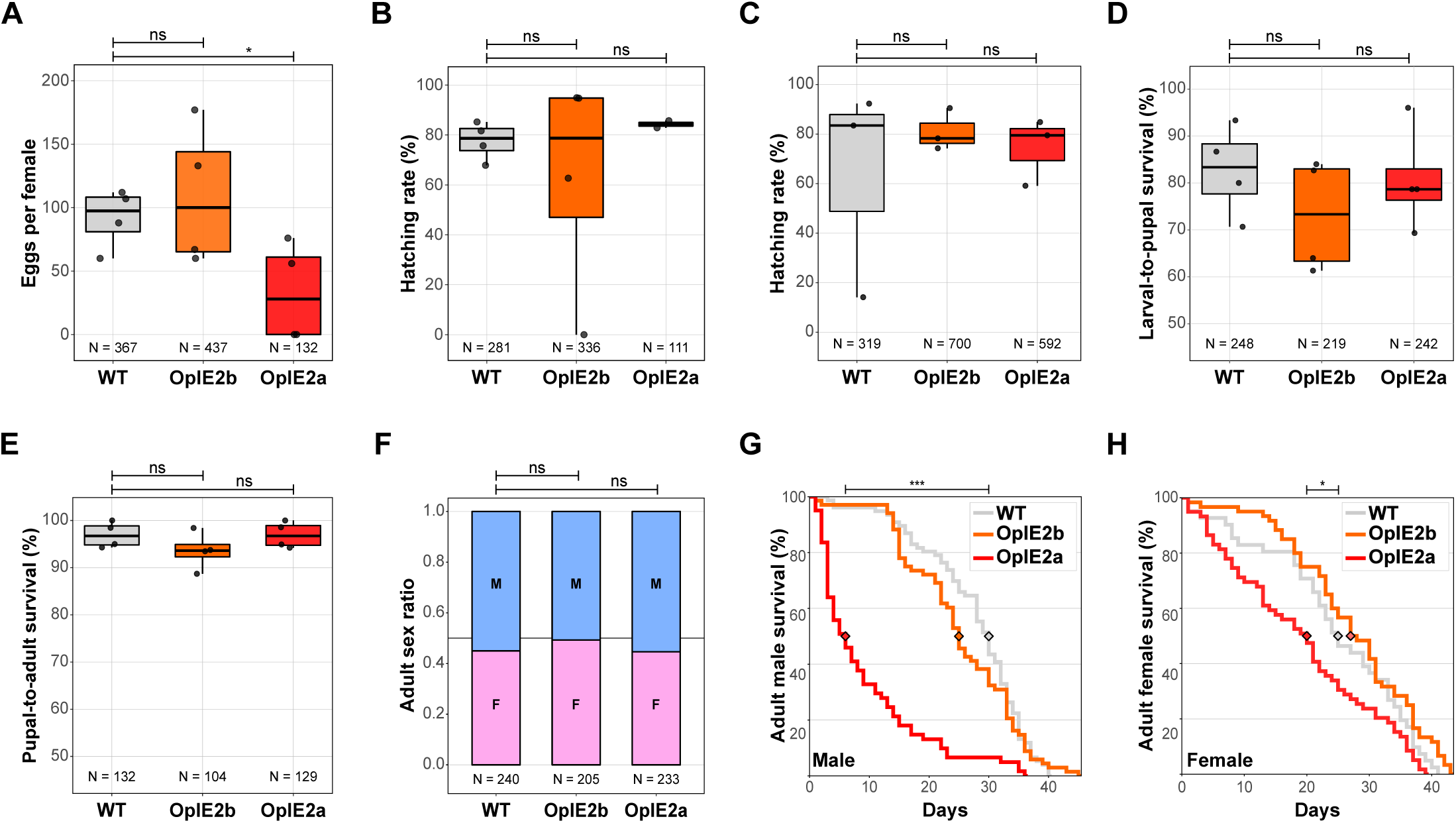
Reproductive output and fitness of *OpIE2*-DsRed strains. **(A–B)** Reproductive performance of females that successfully blood-fed. **(A)** Number of eggs laid per female and **(B)** corresponding hatching rate (% L1 per egg batch). Each point represents an individual female; Boxplots show medians ± IQR; N indicates the total number of eggs and hatched larvae tested per female. Statistical significance was derived from unpaired two-sided *t*-tests, assuming equal or unequal variances, comparing the feeding rates between the different strains to the wild-type control (αL=L0.05). *OpIE2a*-DsRed females produced significantly fewer eggs than wild-type (WT) or *OpIE2b*-DsRed females, while hatching rates did not differ significantly. **(C–F)** Fitness parameters from crosses of transgenic males to WT females. **(C)** Hatching rate (% L1 per egg batch), **(D)** larval-to-pupal survival (%) and **(E)** pupal-to-adult survival (%) were comparable across strains. Boxplots show medians ± IQR; points represent replicate cages. **(F)** Adult sex ratios of progeny from the same crosses. Stacked bars show the mean proportion of males and females. Statistical comparisons were calculated relative to WT control using unpaired two-sided *t*-tests; no significant differences were detected (all *p* > 0.05). **(G–H)** Adult longevity of males **(G)** and females **(H)**. *OpIE2a*-DsRed males exhibited a markedly reduced lifespan compared to WT and *OpIE2b*-DsRed males, while females showed a moderate but significant reduction in survival. Survival curves were analyzed using Kaplan-Meier estimates and compared using log-rank statistics. Diamonds denote median survival per strain.

To evaluate potential effects on male fitness, we crossed transgenic males from each strain to wild-type females, with wild-type intercrosses as controls. Egg hatching, larval-to-pupal, and pupal-to-adult survival rates were comparable across all groups (unpaired two-sided *t*-tests, all *p* > 0.05; **Figure 4C–E**), and adult sex ratios did not differ significantly (**Figure 4F**). However, adult longevity was severely reduced in *OpIE2a*-DsRed males, which had a median lifespan of only 6 days compared to 30 days in wild-type males (log-rank test, *p* < 0.001; **Figure 4G**). *OpIE2b*-DsRed males showed no difference in survival relative to controls (median 25 days vs 30 days, *p* = 0.355). Among females, *OpIE2a*-DsRed also exhibited a modest reduction in survival (median 20 days vs 25 days in wild type, *p* = 0.035), whereas *OpIE2b*-DsRed females survived comparably to controls (median 28 days vs 25 days, *p* = 0.165, **Figure 4H**). Together, these results overturned our initial conclusion that the *OpIE2a*-DsRed marker negatively affected fitness in a female-specific manner, by being detrimental to adult longevity, particularly among males, whereas the *OpIE2b*-DsRed variant maintains normal fecundity, survival, and developmental progression.

## Discussion

In this study, we demonstrate that expression of the fluorescent marker DsRed under the control of a specific variant of the *OpIE2* promoter (*OpIE2a*) causes profound and unexpected phenotypic effects in *An. gambiae*. Females carrying the *OpIE2a*-DsRed marker were unable to blood-feed and were therefore sterile, while males exhibited markedly reduced adult lifespan. In contrast, an alternative promoter variant, which we called *OpIE2b*, drove robust marker expression without measurable effects on feeding, development or longevity. These results reveal that even minor differences in promoter architecture can produce major biological consequences, emphasizing the need to validate commonly used regulatory elements when deployed in new species or experimental contexts.

The female blood-feeding impairment associated with *OpIE2a* was fully penetrant and heritable, persisting across generations - some lines are now over 30 generations old. Behavioral analyses showed that females were normally attracted to the feeder and initiated probing, but were unable to transition to sustained feeding. Microscopic examination identified a fully penetrant curvature of the proboscis and maxillary palps in *OpIE2a* adults of both sexes, suggesting a structural rather than sensory or motivational basis for the defect. This distal bend likely prevents proper fascicle unsheathing and membrane penetration of the proboscis (Jones 1978; Kong and Wu 2010), explaining the repeated probing-grooming cycles and failure to feed. In males, the same curvature may interfere with sugar uptake, consistent with their severely shortened lifespan.

The difference between the *OpIE2a* and *OpIE2b* promoters lies in a 66-bp extension of *OpIE2a* into the viral *IE-2* coding region, which adds 22 amino acids fused to the N-terminus of DsRed. Although both variants contain the core regulatory motifs required for promoter activity (Theilmann and Stewart 1992), this fusion likely alters the transcriptional or translation context, potentially modifying tissue specificity or protein localization. The presence of DsRed fluorescence within the proboscis and maxillary palps of *OpIE2a* mosquitoes supports this hypothesis and suggests that local expression or toxicity in mouthparts underlies the morphological phenotype. While we did not directly measure promoter activity in dissected tissues or test other reporters under the same sequence, the data clearly demonstrate that even seemingly small differences in cloned promoter sequences can markedly change expression domains in transgenic insects, as we have previously observed a native germline promoter in transgenic *An. gambiae* strains (Papathanos et al. 2009).

Beyond its mechanistic implications, this finding carries practical significance for transgenic mosquito research and genetic control tool development. First, transgenic reporters are typically assumed to be phenotypically neutral (Catteruccia, Godfray, and Crisanti 2003; McArthur, Meredith, and Eggleston 2014); however as we have shown here, this assumption needs to be confirmed independently each time, since even standard markers can impose unexpected fitness costs when newly introduced into an engineered insect (Irvin et al. 2004; Marrelli et al. 2006).

In the case of *OpIE2a*-DsRed, the apparent female-specific sterility initially seemed attractive for engineering dominant female lethality, for example in self-limiting or split gene-drive systems (Champer et al. 2022; Verkuijl et al. 2025). However, the severe reduction in male longevity makes this configuration unsuitable for practical applications of population-suppression gene drives in its current form. If expression could be restricted to females, for example through the use of sex-specific splicing (Fu et al. 2010; Liu, Rayes, and Akbari 2024; Scali et al. 2005; Spinner et al. 2022; Weng et al. 2023, 2024), the dominant appendages phenotype could be repurposed as a female-specific lethal trait in future designs.

Finally, the identification of *OpIE2b*-DsRed as a functionally neutral marker with bright, broad expression provides a valuable addition to the transgenesis toolkit available for *An. gambiae*. Because its expression pattern is distinct from commonly used markers such as *3xP3* or *Actin5C*, *OpIE2b* is well suited for multiplexed or combinatorial transgenic designs, including split gene drives and multi-color labeling systems.

In conclusion, our work reveals that promoter sequences alone can profoundly influence expression and phenotype in transgenic insects, transforming an otherwise inert fluorescent reporter into a dominant morphological impairment of mosquito physiology. This work offers both a cautionary example for construct design and a validated promoter variant (*OpIE2b*) suitable for future *An. gambiae* genetic engineering and gene-drive applications.

## Materials and methods

### Mosquito rearing

*An. gambiae* mosquitoes from the G3 (wild-type) strain, and all transgenic mosquito lines were reared under standard conditions of 28°C +/− 2°C and 80% +/− 10% relative humidity. Larvae were fed Essence fish food (Alltech Coppens), while adult mosquitoes were provided with a 10% (wt/vol) sucrose solution. For egg production, adult mosquitoes were allowed to mate for 5-7 days and then fed bovine blood using an artificial feeder (Hemotek, Ltd). An overnight egg-bowl was placed in the mosquito cages 48 hours post-blood-feeding for oviposition. Eggs were allowed 48 hours to hatch, and first instar larvae (L1) were transferred from the egg-bowl to plastic trays, each containing approximately 250 larvae. The larvae were provided with ground fish food daily until they reached the pupal stage.

### Plasmid construction

The *OpIE2-*DsRed transformation constructs were assembled using Golden-Gate assembly. Both plasmids contained an *OpIE2* promoter driving DsRed expression, differing only in the promoter variant used. The *OpIE2a* promoter fragment (541 bp) and the *OpIE2b* promoter fragment (652 bp) were PCR-amplified with primers incorporating BsaI recognition sites and compatible overhangs for directional assembly. Each promoter fragment was assembled upstream of the DsRed coding sequence to generate an *OpIE2*-DsRed cassette. In both constructs, the cassette was flanked by *piggyBac* inverted terminal repeats and a single *attB* site, enabling either random *piggyBac*-mediated or *phiC31* site-specific integration. Final plasmids were validated by Oxford Nanopore sequencing (Plasmidsaurus) to confirm promoter sequence identity and integrity of the assembled cassette.

### Embryo microinjections

Microinjections were performed into eggs of the *attP*-E strain, a homozygous transgenic line carrying an *attP* docking site on chromosome 3 and a *3xP3*-CFP fluorescence marker. *An. gambiae attP*-E eggs were gently aligned on a glass slide against a strip of thin nitrocellulose membrane, with moisture maintained using UltraPure™ Distilled Water (Invitrogen) applied to dampened filter paper. Using a fine brush, eggs were oriented with the posterior end facing upwards to maximize the chances of germline integration. A mixture of 300 ng/µl of the transformation construct and 100 ng/µl of a helper construct encoding vasa-driven *phiC31* integrase was injected. Hatched larvae were screened for transient expression of the *OpIE2*-DsRed marker, and all transient-positive and transgenic individuals were backcrossed to G3 wild-type mosquitoes. Progeny were maintained under standard rearing conditions.

### Blood-feeding assay and recording of female behavior

Adult males and females were allowed to mate for 5–7 days prior to blood-feeding. For each strain, three biological replicates of 14–20 females were transferred to Bugdorm minicages (Model BD4S1515) for feeding. Females were offered blood for 25 min using the Hemotek Ltd membrane feeding system, with reservoirs filled with thawed bovine blood and sealed with stretched Parafilm. Feeding sessions were recorded using an Arducam camera system with underlighting to document female behavior for subsequent analysis. Reservoirs were replaced with freshly thawed blood and new Parafilm membranes between assays performed on different strains. If no feeding was observed, a 15 min rest period was followed by a stimulation feeding session of 25 min, using the same Hemotek reservoir. During stimulation, air was exhaled intermittently onto the reservoir surface to increase female attraction. At the end of each assay, females were anesthetized at 4°C, and those with visibly blood-filled abdomens were aspirated and counted as successfully fed. Blood-feeding recordings were analyzed using a trained behavioral detection and classification neural network built on the YOLOv11 object detector (Khanam and Hussain 2024), which was fine-tuned for mosquito detection and adapted from the behavioral analysis pipeline developed by Vainer et al. (2024). The pre-trained YOLOv11l model was tuned using a curated dataset of annotated images. Detection parameters were set to a confidence threshold of 0.2 and an intersection-over-union (IoU) threshold of 0.5 for bounding-box predictions. Feeder occupancy was analyzed using a linear mixed model with the different strains and time as fixed effects and replicate as a random intercept.

Blood-feeding videos recorded during the assay were additionally analyzed using the open-source event-logging software BORIS - Behavioral Observation Research Interactive Software (Friard and Gamba 2016). For each strain, one video with the highest number of females was selected for analysis (G3: 20 females, replicate R1; OpIE2b: 20 females, replicate R3; OpIE2a: 20 females, replicate R1; see **Supplementary Table 3**). Female behavior was tracked for the first 5 min following placement of the Hemotek feeder in the cage (time = 0). Because individual females could not be distinguished, behaviors were scored per visit to the feeder area. A visit was defined as entry into the area of interest over the Hemotek feeder, with multiple visits by the same female possible. For each visit, arrival and departure times, the number of probing events and, where applicable, the start and end times of blood-feeding were recorded.

### Female fertility assays

Females that blood-fed during the assay were individually isolated for oviposition. The number of females per strain was determined by the number of *OpIE2a*-DsRed females that successfully fed after stimulation. Wild-type and *OpIE2b*-DsRed females were randomly selected to match this number. 48 h post blood-feeding, an egg-bowl was placed in each cage and removed the following day. Egg papers were photographed for counting using Clickmaster2000 (https://github.com/EuracBiomedicalResearch/clickmaster2000). Larvae were provided food 24 h after egg collection, and the number of hatched larvae was recorded 24 h later to calculate hatching rates per female. Progeny from these assays were subsequently reared to adulthood and tested for blood-feeding behavior as described above (filming not performed due to technical limitations).

### Fitness and survival assays

To evaluate potential fitness costs of the *OpIE2*-DsRed cassette, paternal crosses were established between transgenic males and wild-type females, with wild-type intercrosses as controls. Eggs were collected and counted to determine hatching rates. Hatchlings were distributed into trays at a density of 75 larvae per tray and reared under standard conditions. At pupation, male and female pupae were collected and counted. Pupae were allowed to emerge in Bugdorm minicages (Model BD4S1515), and the number of adults was determined by counting remaining pupal cases. These data were used to calculate larval-to-pupal and pupal-to-adult survival rates for each sex and strain, as well as the sex ratio of emerging adults. Adult longevity was assessed in Bugdorm minicages (Model BD4S1515) containing 30–35 adults (24–72 h post-emergence) per strain, maintained on 10% sucrose solution. Dead adults were recorded and removed daily until all individuals had died. If sex or age at death could not be determined, the individual was censored on the day of discovery. Sucrose solution was replaced with a new, clean feeder every 14 days. Survival data were analyzed using Kaplan– Meier survival curves with log-rank tests.

### Mosquito imaging

Photos of larvae, pupae, and adults were captured under standardized conditions to ensure consistent exposure across strains. Larvae at the fourth instar and pupae were imaged using an Olympus SZX16 binocular equipped with a U-HGLGPS fluorescence illumination system. All fluorescence images were acquired using identical exposure settings for each strain. Adult mosquitoes were imaged using a Nikon SMZ25 stereo microscope. 2 to 3 day old adults were first anesthetized with CO₂, after which legs and wings were removed to expose the body and proboscis for imaging. Dissected adults were then transferred to 50 mL plastic cups sealed with Parafilm containing silica gel desiccant to prevent condensation and placed at −20 °C for 30 min to achieve full immobilization. Following freezing, cups were thawed briefly at room temperature immediately before imaging. Fluorescence parameters and exposure settings were kept identical between strains to allow direct comparison of DsRed signal patterns.

### Statistical analysis

Statistical tests were performed in Python using standard packages (3.10.9; scipy 1.15.3, statsmodels 0.14.4) and JMP Pro 18.0.2 software. All experiments were independently replicated a minimum of three times unless stated otherwise, and each replicate represented an independent biological group (e.g., cage or clutch). The number of individuals or replicates used for each analysis is indicated in the figure panels and methods sections.

## Author contributions

AS and PAP conceived and designed the study. AS, ESY, ESMK and LBL generated transgenic mosquito lines. AS performed behavioral, imaging, and fitness assays, and analyzed the data. ARL helped with figure illustrations. ES-S and JB developed the trained behavioral detection and classification neural network. ESMK, ESY, and LBL assisted with mosquito genetic engineering. TP and GP contributed to adult proboscis imaging. PAP supervised the project, secured funding, and wrote the manuscript together with AS with input from all authors. All authors read and approved the final manuscript.

## Acknowledgements

We would like to thank Daniella An Haber and Yael Arien for their valuable discussions and critical feedback and Gleb Ens, Ruth Shacham and Albert Nazarov for technical support.

## Ethics

All animals were handled in accordance and under the supervision of the ARO Institutional Animal Care and Use Committee approval number 2307-118-2-VOL-IL. All insect work was performed in facilities maintaining Arthropod Containment Level 2. This work received Institutional Approval and relevant authorizations from the Israel Ministry of Environmental Protection and Ministry of Agriculture (#31/2019).

---

**Supplementary Figure 1. Quantification of *OpIE2*-DsRed female feeding behavior. (A)** Time spent on the feeder (min) per female visit. The time that each female spent on the feeder area differed significantly between strains compared to the WT control (Kruskal–Wallis, *H* = 37.6634, *p* < 0.001). *OpIE2b*-DsRed females spent slightly more time on the feeder compared to WT females (Dunn’s comparisons with Bonferroni correction vs WT, *p* = 0.033), whereas *OpIE2a*-DsRed females spent significantly less time (*p* < 0.001). **(B)** Frequency of probing events per female visit. Group differences were significant (Kruskal–Wallis, *H* = 9.93, *p* = 0.007). *OpIE2b*-DsRed females showed a slight reduction in probing frequency compared to wild-type females (Dunn’s comparisons with Bonferroni correction vs WT, *p* = 0.0485), while *OpIE2a*-DsRed females did not differ significantly (*p* = 0.85). **(C)** Frequency of proboscis grooming events per female visit. Group differences were significant (Kruskal–Wallis, *H* = 11.24, *p* = 0.0036). Grooming frequency was significantly higher in *OpIE2a*-DsRed females relative to WT (Dunn’s comparisons with Bonferroni correction vs WT, *p* = 0.042), whereas *OpIE2b*-DsRed females were not different from WT (*p* = 0.66). For WT and *OpIE2b*-DsRed females, probing and proboscis grooming events prior to the initiation of blood-feeding are considered, to enable comparison with *OpIE2a*-DsRed females, which do not initiate feeding.

**Supplementary Figure 2. Quantification of proboscis curvature in OpIE2-DsRed mosquitoes. (A)** Mean ventral curvature angle angle (°) of female proboscises. *OpIE2a*-DsRed females exhibited a significantly greater ventral curvature relative to wild-type females (unpaired two-sided *t*-test, *p* = 0.007), whereas OpIE2b-DsRed females did not differ significantly from controls (*p* = 0.63). Each data point represents an individual mosquito. **(B)** Mean ventral curvature angle (°) of male proboscises. *OpIE2a*-DsRed males exhibited a significantly greater ventral curvature relative to wild-type males (unpaired two-sided *t*-test, *p* < 0.001. No significant difference was observed between *OpIE2b*-DsRed and wild-type males (*p* = 0.43).

**Supplementary Table 1.**
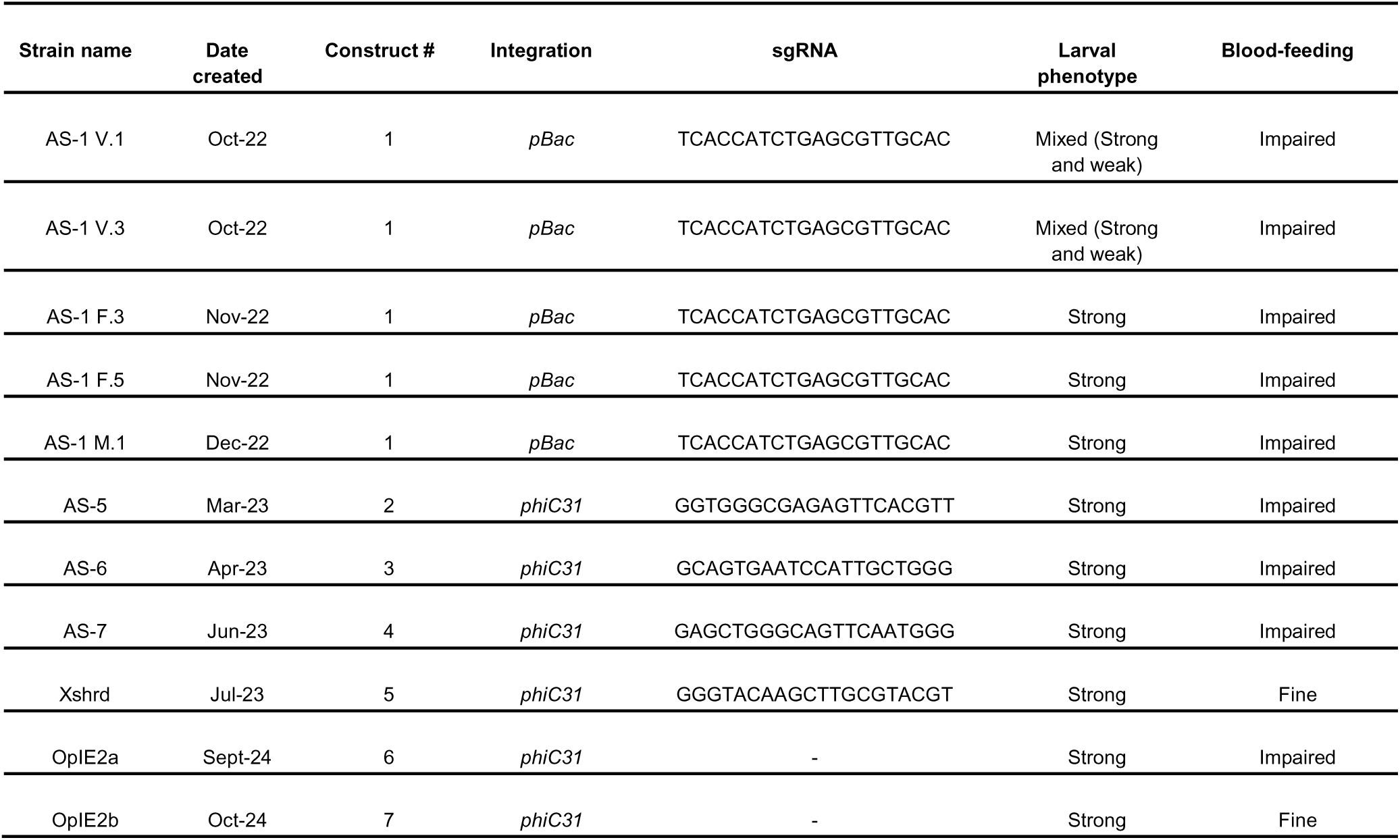
Summary of transgenic strains. Each strain is identified by an internal strain name. Construct number refers to the numbering system used to distinguish transformation plasmids. Integration method indicates whether transformation occurred through random *piggyBac*-mediated transposition or *phiC31*-mediated site-specific integration. The sgRNA linker sequence shows the 5′-3′ sgRNA space sequence cloned for each construct containing a *U6*-sgRNA cassette. Larval marker phenotype denotes the strength and consistency of DsRed fluorescence, while female blood-feeding behavior and male fertility summarize adult phenotypic outcomes for each line.

**Supplementary Table 2.** Heritable disruption of blood-feeding by *Opie2a-*DsRed. Daughters of the two *OpIE2a*-DsRed females that successfully fed after additional stimulation (Figure 2C) were tested for blood-feeding using the same experimental setup as their mothers. Three replicate cages were established, each containing nine 5–7-day-old females previously mated to wild-type males. In all replicates, no females were observed with visibly blood-filled abdomens at the end of the feeding period, confirming complete penetrance and transmission of the non-feeding phenotype.

| Strain | Experiment date | Repeat | N females | N females fed |
| --- | --- | --- | --- | --- |
| OpIE2a females | 05.01.2024 | 1 | 9 | 0 |
| OpIE2a females | 05.01.2024 | 2 | 9 | 0 |
| OpIE2a females | 05.01.2024 | 3 | 9 | 0 |

**Supplementary File 1. Sequences of the *OpIE2a* and *OpIE2b* promoter variants and constructs generated in this study in FASTA format.** File numbering corresponds to the constructs listed in **Supplementary Table 1** and shown in **Figure 1A**.

**Supplementary File 2. Manual annotations of behavioral metrics of membrane feeding assays.** The table contains output data of BORIS manually recorded behavioral observations for individual *An. gambiae* females of the indicated genotypes. Each row represents a single female visit. Behavioral events were recorded from 5-min video segments and include the number of grooming and probing events, total time spent in the area of interest (AOI) near the feeder, and total duration of blood-feeding. “Time at AOI” represents cumulative occupancy on the feeder, while “ROI before feeding” and associated probing metrics quantify pre-feeding activity. Frequencies are expressed as events per second or minute as indicated.

**Supplementary File 3. Statistical analysis summary.** All statistical tests, *p*-values, and significance levels corresponding to figures and supplementary figures presented in this study.

