## Supplementary figures and images for "An *OpIE2*-DsRed marker disrupts female blood-feeding and shortens lifespan in the malaria vector *Anopheles gambiae*"

### Supplementary Figures

#### Supplementary Figure 1

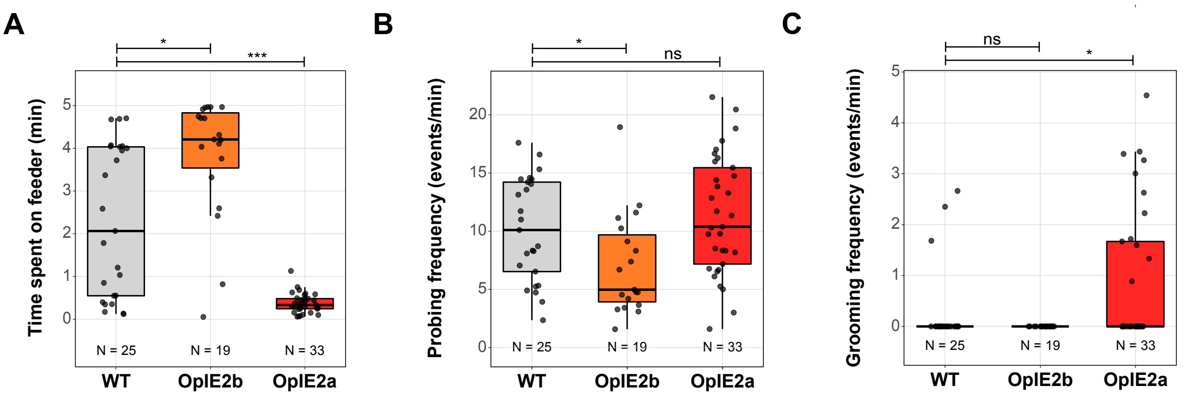


#### Supplementary Figure 2

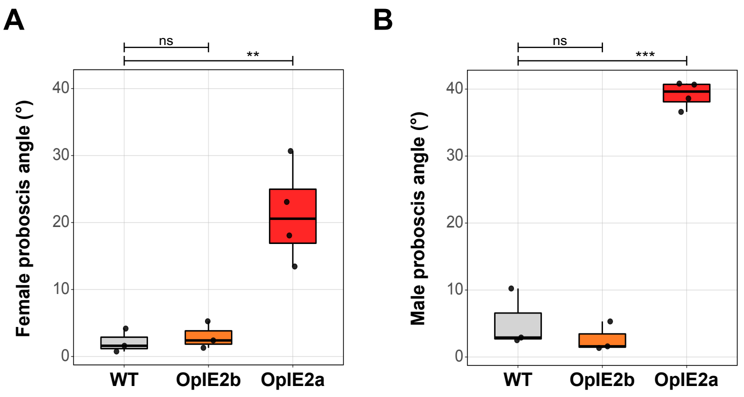
